# The soil microbial community alters patterns of selection on flowering time and fitness related traits in *Ipomoea purpurea*

**DOI:** 10.1101/672691

**Authors:** Lindsay Chaney, Regina S. Baucom

## Abstract

**Premise of the study:** Plant flowering time plays an important role in plant fitness and thus evolutionary processes. Soil microbial communities are diverse and have a large impact, both positive and negative, on the host plant. However, owing to few available studies, how the soil microbial community may influence the evolutionary response of plant populations is not well understood. Here we sought to uncover if below-ground microbial communities act as an agent of selection on flowering and growth traits in the common morning glory, *Ipomoea purpurea*.

**Methods:** We performed a controlled greenhouse experiment in which genetic lines of *I. purpurea* were planted into either sterilized soils, or soils that were sterilized and re-inoculated with the microbial community from original field soil. This allowed us to directly test the influence of alterations to the microbial community on plant growth, flowering, and fitness, as well as assess patterns of selection in both soil microbial environments.

**Results:** We found that a more complex soil microbial community resulted in larger plants that produced more flowers. Selection strongly favored earlier flowering when plants were grown in the complex microbial environment than compared to sterilized soil. Additionally, we uncovered a pattern of negative correlational selection on growth rate and flowering time, indicating that selection favored different combinations of growth and flowering traits in the simplified versus complex soil community.

**Conclusions:** Together these results suggest the soil microbial community is a selective agent on flowering time and ultimately that soil microbial community influences important plant evolutionary processes.

## INTRODUCTION

Plant flowering time plays an important role in plant fitness and thus evolutionary processes. For example, a plant that flowers too early or too late in the season may be misaligned with pollinators, decreasing gene flow and potentially fitness through pollen limitation or inbreeding depression (Elzinga et al., 2007; Franks, 2015). Likewise, early flowering is associated with greater susceptibility to damage from herbivory or disease, ultimately leading to lower fitness (Inouye, 2008). The timing of flowering is itself impacted by various abiotic factors – temperature, drought, photoperiod, soil nutrients, and ambient CO_2_ (*reviewed in* Kazan and Lyons, 2016; Blackman, 2017; Cho et al., 2017) – as well as biotic factors such as plant pathogens, herbivory, and soil microbes (Lau and Lennon, 2011, 2012; Züst et al., 2011; Panke-Buisse et al., 2014; Wagner et al., 2014; Lyons et al., 2015). Of these factors, the influence of the soil microbial community on flowering time has come under increasing scrutiny given the recent recognition of the importance of plant-soil feedbacks on fitness (Bever, 2003; Reynolds et al., 2003; Geml and Wagner, 2018). Understanding how the soil microbial community alters flowering phenology is thus of broad interest to researchers aiming to optimize crop yield (Gopal and Gupta, 2016), as well as those working on plant eco-evolutionary feedbacks (terHorst and Zee, 2016), adaptation to novel environments (Lau and Lennon, 2012), and climate change (Classen et al., 2015).

Despite the increasing interest in the role of soil microbes on plant phenological patterns, only a handful of studies have examined their influence by performing experimental manipulations of the overall soil microbial community. Results of this work appear to be somewhat mixed: while modification of the soil microbial community led to both earlier and later flowering time in *Brassica rapa* and the wild mustard *Boechera stricta* (Panke-Buisse et al., 2014; Wagner et al., 2014), a simplified soil microbial community, compared to more diverse soils, did not alter flowering time in *B. rapa* (Lau and Lennon, 2011). Strikingly, the majority of studies investigating the role of the soil microbial community on flowering phenology have focused largely on close relatives of *Arabidopsis* (Lau and Lennon, 2011; Panke-Buisse et al., 2014; Wagner et al., 2014), meaning that our current understanding of the effects of the soil microbial community on flowering time and fitness are somewhat taxonomically limited.

However, a consistent finding across studies is that soil microbial communities can alter patterns of selection on flowering time, and in this way can impact plant evolutionary processes. In simplified soil communities, *B. rapa* individuals that flowered earlier were at a fitness advantage compared to those that flowered later, an effect that was stronger than the abiotic stress of low soil moisture (Lau and Lennon, 2011). Exposure to different natural soil microbiota likewise altered patterns of selection on flowering time in *B. stricta* – both increasing and decreasing fitness depending on the soil type (Wagner et al., 2014). Interestingly, these same natural soils, when sterilized, did not lead to differential patterns of selection, indicating that chemical differences in the soils did not differentially influence fitness as did the presence of the microbial community. The results from *B. rapa* and *B. stricta* together show that the soil microbial community is an important agent of selection on flowering time – perhaps even more so than that of abiotic stressors. As above, however, this work was performed using close relatives of *Arabidopsis*. Can we expect that the soil microbial community acts as an agent of selection on flowering time in other species?

Additionally, the soil microbial community is associated with altering plant growth and size across species (Lugtenberg and Kamilova, 2009; Henning et al., 2016), and similar to flowering time, there is evidence that the soil microbial community can act as an agent of selection for increased plant biomass (Lau and Lennon, 2011). If and how soil microbes influence the optimal transition between growth and flowering, however, has yet to be examined in any species. The life-history switch between growth and flowering is complex and modulated by factors that are intrinsic to the plant – age and hormone levels – as well as extrinsic environmental cues, such as photoperiod, temperature, light intensity and spectral quality (Kazan and Lyons, 2016; Blackman, 2017; Cho et al., 2017). Given the importance of the transition between growth and flowering, and its environmental dependency, selection should favor a transition in life-history stage that will optimize fitness in any particular environment. One could expect, for example, that fitness would be highest in plants that grow at an optimal rate to support the onset of flowering that best aligned with pollinators or simply the production of fruits. Evidence for selection on an optimal plant growth/flowering time combination would be provided by a pattern of correlational selection between plant growth traits and flowering time in a selection analysis; an altered pattern of correlational selection between plant growth and flowering – following changes in the soil microbial community – would provide evidence that the soil community imposes selection on this important life history transition.

In this study, we examine the potential that the microbial soil community alters growth and flowering and acts as an agent of selection on these traits in *Ipomoea purpurea* (Convolvulaceae), the common morning glory. *I. purpurea* is a weedy annual vine most often found in agricultural areas or areas of high soil disturbance. It is an appropriate model species for addressing the influence of soil microbes on flowering time since it is able to thrive across a variety of soil types in disparately located natural populations, it exhibits rapid growth to flowering (seed to flowering within three weeks), and produces a relatively large number of flowers in greenhouse conditions (personal observations, RS Baucom). Additionally, the species shows a high correlation between flower number and seed set in field conditions (r = 0.92), allowing us to use flower number (reproductive output) as a proxy for overall fitness. To test the impact of the soil microbial community on flowering time, we imposed treatments that altered the diversity of soil microbial community in a greenhouse setting, and recorded flowering time, plant size and growth, and reproductive output to address three main questions: 1) Are plant growth and fitness traits influenced by soil microbial environment? 2) What are the patterns of selection on these traits?, and 3) Do patterns of selection change as result of soil microbial environment?

We addressed these questions by planting replicate seeds from inbred maternal lines of *I. purpurea* in simplified and complex soils, allowing us to test for the presence of genetic variation underlying flowering phenology and growth traits (genotypic variation), the influence of altering the microbial community on patterns of plant flowering (variation due to microbial environment), and the potential that genotypes responded differently to the altered soil microbial communities (adaptive plasticity, or GXE). Our results support the idea that the soil microbial community influences fitness of *I. purpurea* individuals, and similar to *B. rapa* and *B. stricta*, acts as an agent of selection on flowering time. We likewise show that the soil microbial community influences the pattern of selection on combinations of growth and flowering. Overall, our findings add to the growing body of knowledge showing that the soil microbial community impacts selection on flowering time, and in this way can influence important plant evolutionary processes.

## MATERIALS AND METHODS

### Experimental Procedure

#### Plant material

We used replicate seeds of 20 inbred maternal lines (hereafter genotype) of *Ipomoea purpurea*, which originated from a population generously supplied by M. Rausher and were collected in Durham, North Carolina in 1985 and inbred in the greenhouse for about 15 generations.

#### Soil treatments

We manipulated the soil microbial community by autoclaving field soil (hereafter simple soil microbial community) and then re-inoculating the sterilized soil using a slurry solution derived from field soil (hereafter complex soil microbial community). To do so, we collected soil for the experiment from an agricultural field at the University of Cincinnati’s Center for Field Studies in Harrison, Ohio (identified as silt loam, pH 7.1; Michigan State University Soil and Plant Nutrient Laboratory, East Lansing, MI). We autoclaved all soil two times at 121°C and 15 PSI for 2 hours each cycle. After autoclaving, we mixed soil with sterile perlite (4:1) to improve drainage and aeration without changing the pH or nutrient content of the soil. For the complex microbial community, 0.5 liters non-autoclaved field soil was soaked with 2 liters of sterile water overnight, centrifuged at 1,000 g, and then the aqueous layer was poured back onto half of the sterilized soil. This process preserved the natural soil microbes and re-introduced them into the autoclaved soil. We modeled our soil preparation on studies showing that autoclaving leads to a simple microbial community, and re-inoculated/autoclaved soils house a more complex microbial community. In this design, any differences between treatments is attributed to the addition of inocula (Marschner and Rumberger, 2004; Berns et al., 2008; Lau and Lennon, 2011; Aschehoug et al., 2012; Panke-Buisse et al., 2014).

#### Experimental design

Six replicate seed from each of the twenty genotypes were planted into the two soil microbial treatments (overall N = 240). To ensure establishment, seeds were first scarified by nicking the seed coat with a razor blade then placed in petri dishes with sterile water under grow-lights. Once emerged, seedlings of each genotype were planted into prepared soil treatments in 4-inch pots. Treatments were planted subsequently to prevent cross-contamination. We then placed pots in a completely randomized design in the University of Cincinnati greenhouse. Plants were grown under metal halide lamps with a daily 12 hr light regime to promote flowering and were watered daily. We treated plants with Blossom Booster (JR Peters INC.; Allentown, PA) every 10 days. The duration of the experiment was 21 weeks, which is approximately the length of time *I. purpurea* persists until senescence in field conditions.

#### Plant phenotypic measurements

We focused on four phenotypic traits: flowering day, plant size, growth, and total number of flowers. Flowering day is a record of the first day a plant began to flower. Plant size was estimated by summing the lengths of all true leaves from each plant 34 days after planting. The sum of leaf lengths was previously found to be an excellent predictor of dry shoot biomass in this species (*R*^2^ = 0.891, Chaney and Baucom, 2014). Plant growth was estimated as the height of the plant 34 days after planting minus the height at 14 days after planting, divided by the number of days elapsed. The total number of flowers is a record of the sum of flowers produced by each plant throughout the experiment; flower number was recorded five days a week for all plants.

### Data analysis

#### Phenotypic response

We used a series of univariate linear mixed-models from the *lme4* package (Bates et al., 2015) to examine the effects of genotype, soil microbial community, and their interaction on plant flowering day, size, growth, and total number of flowers. For each model, the response variable was one of the four phenotypic traits, soil microbe treatment was a fixed effect; the genotype (maternal line) and the interaction of genotype by treatment were random effects. The significance of effects was determined using the *lmerTest* package (Kuznetsova et al., 2017). The residuals of flowering day were left-skewed, so a square-root transformation was performed to meet assumptions of the model. All other response variables met the assumptions of normality and equal variance.

#### Phenotypic selection

To determine if the soil microbial community is a selective agent on flowering time, size, and growth, we used phenotypic selection analysis (Lande and Arnold, 1983) to estimate both selection differentials and selection gradients on each trait. The total number of flowers was used as a proxy for fitness in all selection analyses and is generally considered ‘reproductive potential’ rather than life-time fitness, but is an excellent predictor of total number of seed (*r* = 0.924, *t* = 13.23, *N* = 32 (maternal lines), *P* < 0.01; R. S. Baucom and R. Mauricio, unpublished data). Relative fitness was calculated for each individual as the total number of flowers divided by the overall mean fitness of the experimental population. Phenotypic traits were standardized to a mean of zero and a variance of one prior to analysis. We estimated selection differentials (*S*) using a univariate regression of relative fitness on each trait separately. This is a measure of the total selection acting on a trait due to both direct and indirect selection. Selection gradients, which measure only direct selection on each trait by accounting for correlations with other traits in the model, were calculated by performing multiple regression of relative fitness on all phenotypic traits together. We estimated linear (directional) selection gradients (β) using models containing only the linear terms whereas we estimated non-linear selection gradients (γ) by doubling quadratic regression coefficients in a full model that contained linear terms, quadratic terms, and cross-product terms of focal traits. Nonlinear selection gradients indicate selection that acts on either the phenotypic variance of a trait (quadratic selection) or the phenotypic covariance between two traits (correlational selection). To determine if the significant negative nonlinear selection demonstrates true stabilizing selection, the spread of relative fitness values was examined for an intermediate optimum across the phenotypic distribution of flowering day (Mitchell-Olds and Shaw, 1987). We visualized fitness surfaces for significant correlational selection by performing a thin-plate spline nonparametric regression approach, using the *Tps* function in the *fields* package (Nychka et al., 2015). Smoothing parameters for each spline where chosen to minimize generalized cross-validation score.

We used an analysis of covariance (ANCOVA) to determine if the soil microbial community altered patterns of selection on plant traits. To do this, we performed a univariate regression of relative fitness on the trait in question, soil microbe treatment, and their interaction. A significant interaction between plant traits and soil microbe treatment would indicate that selection differentials differed between treatments. We similarly tested for differences in linear and quadratic selection gradients using a multivariate regression of relative fitness on all plant traits, soil microbe treatment, and their interactions. Differences in selection gradients by soil microbe treatment would be signified by a significant trait and soil treatment interaction. All selection differentials and selection gradients were calculated within each treatment, as stated previously, using only data for that treatment. All analyses were performed in the R statistical environment (v 3.2.1; R Team, 2015). Analysis code and data will be made available via Dryad post-acceptance.

## RESULTS

### Phenotypic differences according to microbial soil environment

Both the size of plants and the total number of flowers plants produced were influenced by differences in soil microbial community (Table 1, Figure 1). We found that plants grown in a complex soil microbial community were 12% larger and had 15% more flowers than plants grown in the simple soil microbial community (*F* = 7.47, *P* = 0.01, and *F* = 4.13, *P* = 0.05; plant size and total number of flowers, respectively). Plants in the complex soil microbial community flowered, on average, a half day later and grew 6% faster (20 mm per day) than plants grown in simple soil microbial community, but the difference between treatments was not significant (*F* = 0.04, *P* = 0.85, and *F* = 1.88, *P* = 0.17; flowering date and growth, respectively). We found evidence of maternal line variation for three of the four traits (flowering day, plant size, and total number of flowers; Table 1); however, there was no indication of genetic variation for plasticity for any of the traits examined (no significant genotype by treatment effect for any trait).

**TABLE 1:** Influence of soil microbial community on *Ipomoea purpurea* plant traits. Trait mean and standard error of the mean in each soil environment. *F*-value and chi-square values showing the effects of genotype, microbe treatments, and interaction on plant phenotype. Significant effects are indicated with asterisks: \**P* < 0.05, \*\**P* < 0.01.

| Trait | Autoclave<br>Inoculum | Autoclave | Genotype | Treatment | Genotype x<br>Treatment |
| --- | --- | --- | --- | --- | --- |
| | mean ± se | mean ± se | $\chi^2$ | *F*-value | $\chi^2$ |
| Flowering Day | 44.89 ± 1.11 | 44.30 ± 1.45 | 9.57** | 0.00 | 0.00 |
| Size | 45.50 ± 1.30 | 40.67 ± 1.30 | 5.50* | 7.47** | 0.00 |
| Growth | 3.63 ± 0.10 | 3.41 ± 0.13 | 0.00 | 1.88 | 0.00 |
| Total Flowers | 67.08 ± 3.56 | 58.16 ± 3.17 | 8.98** | 4.13* | 0.43 |

**FIGURE 1:**
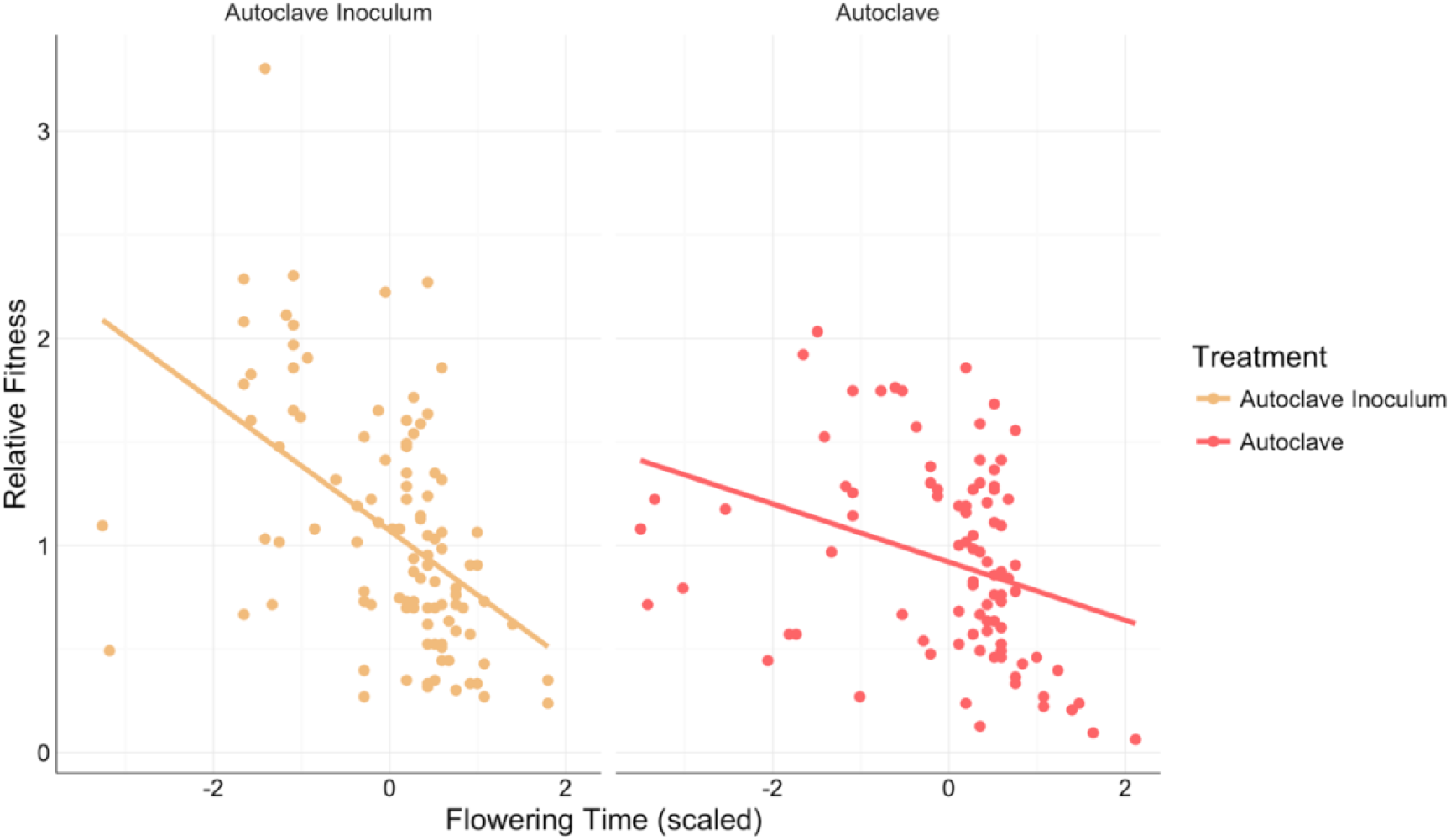
Influence on soil microbial community on flowering day, growth, size, and total number of flowers in *I. purpurea*. Median, upper and lower quantile, upper and lower whiskers, and outliers for each of the two soil treatments is shown.

### Selection differentials

Selection acted in favor of earlier flowering and bigger plant size (Appendix S1). Further, we found a significant interaction between flowering day and soil microbe treatment, indicating that soil microbial community significantly influenced patterns of selection on this trait (Table 2). Specifically, selection on flowering day was stronger in the complex soil microbial community compared to the simple soil microbial community (Figure 2; *F* = 6.04, *P* = 0.02). We likewise uncovered stronger selection for larger plant size in the complex soil microbial community, but this did not significantly differ from the simple soil microbial community (*F* = 1.32, *P* = 0.25).

**TABLE 2:** Influence of soil microbial community on selection differentials. Shown are selection (*S*) values in each treatment. F-value for treatment from the ANCOVA analysis testing the effect of soil microbe treatment on selection differentials. Significant effects are indicated with asterisks: \**P* < 0.05, \*\**P* < 0.01.

|  | Autoclave Inoculum | Autoclave | *F*-value |
| --- | --- | --- | --- |
| Flowering day | -0.31** | -0.14** | 6.04* |
| Size | 0.17** | 0.08 | 1.32 |
| Growth | -0.02 | -0.04 | 0.06 |

**FIGURE 2:**
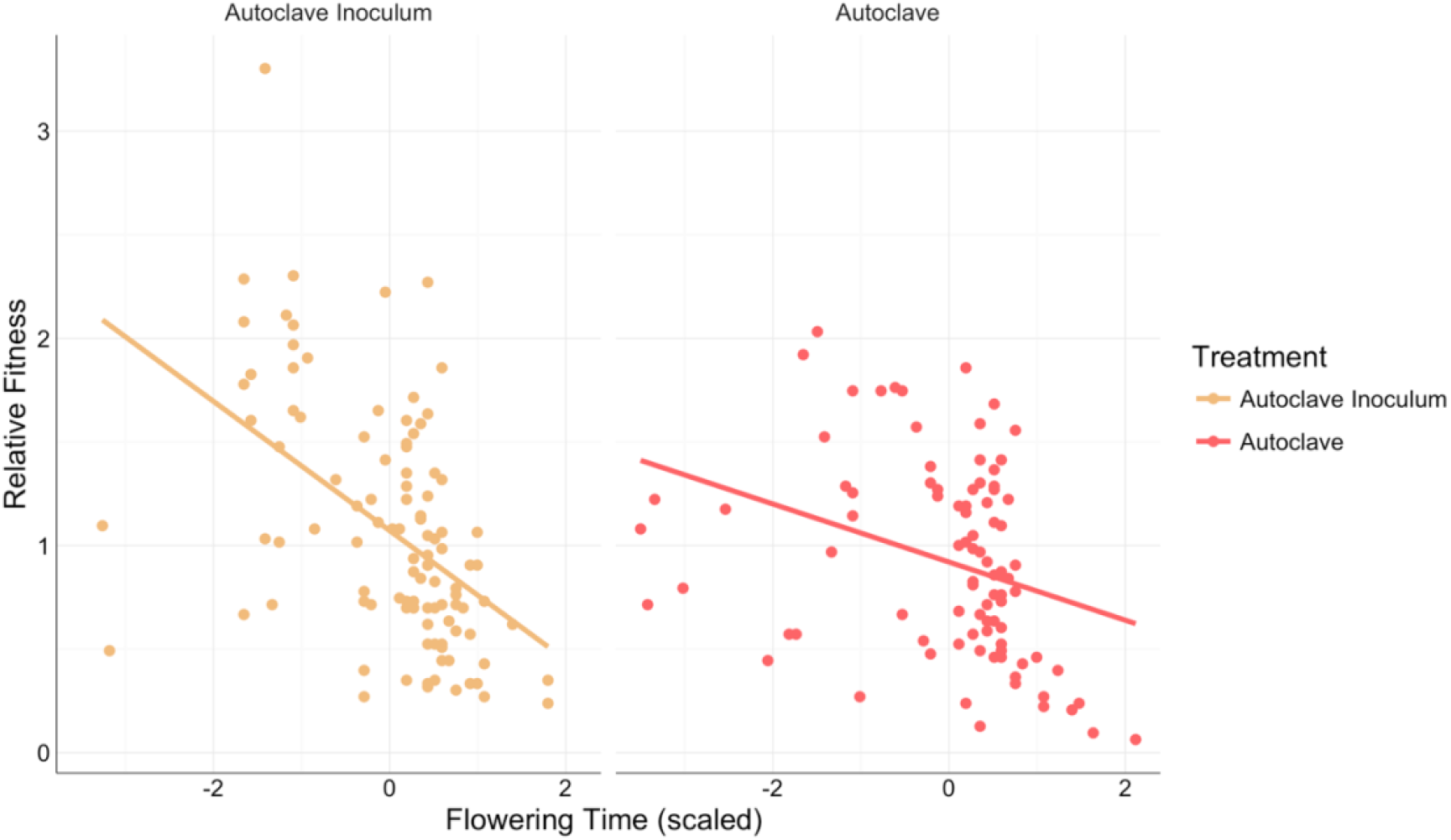
Total selection (S) on *I. purpurea* flowering time in a complex (autoclave + inoculum) and simple (autoclave) soil microbial community.

### Selection gradients

Selection gradients were similar to selection differentials; we found evidence of linear selection favoring earlier flowering and larger plant size (Appendix S2) in both soil microbial environments. We also uncovered evidence of significant non-linear, negative selection on flowering day. Examination of fitness values across the phenotypic distribution of flowering day showed an intermediate fitness optimum, suggesting mid- to early-flowering plants are favored in both soil microbial environments (*i.e.*, significant stabilizing selection). Similar to the analysis of selection differentials, we found that changes to the microbial community significantly affect linear selection on flowering day, with stronger selection occurring in the complex soil microbial community (Table 3; *F* = 5.39, *P* = 0.02). The soil microbial community also differentially influenced patterns of correlational selection on flowering day and growth rate (Figure 3; *F* = 7.82, *P* = 0.01). In the complex soil microbial environment, we identified a positive (but non-significant) interaction between growth rate and flowering (γ = 0.23, P > 0.05; Table 3) whereas the interaction between growth rate and flowering time in the simple soil microbial environment was negative (γ = −0.31, P < 0.05; Table 3). This change in the pattern of selection between soil environments suggests that an optimal combination between growth and flowering (*i.e.*, moderate growth and a mid- to early flowering time) was favored in the complex soil microbial community, whereas two strategies were favored in the simplified soil environment – fast growth and early flowering, or slow growth and late flowering.

**TABLE 3:** Influence of soil microbial community of selection gradients (multivariate selection). Shown are linear (beta; β) and quadratic (gamma; γ) values in each treatment. *F*-value from the ANCOVA analysis testing the effect of soil microbe treatment on selection gradients. Near significant, and significant effects are indicated with asterisks: \**P* < 0.05, \*\**P* < 0.01, \*\*\**P* < 0.001.

| Linear Selection ($\beta$) | Autoclave Inoculum | Autoclave | Treatment |
| --- | --- | --- | --- |
|  |  |  | *F*-value |
| Flowering day | -0.28*** | -0.13** | 5.39* |
| Size | 0.18*** | 0.14* | 0.53 |
| Growth | -0.084 | -0.10 | 0.03 |
| Quadratic selection ($\gamma$) | Autoclave Inoculum | Autoclave | Treatment |
|  |  |  | *F*-value |
| Flowering day<sup>2</sup> | -0.28*** | -0.17** | 0.74 |
| Size<sup>2</sup> | -0.05 | -0.10 | 0.10 |
| Growth<sup>2</sup> | -0.12 | 0.07 | 2.38 |
| Flowering day x Size | -0.21 | 0.21 | 0.88 |
| Flowering day x Growth | 0.23 | -0.30** | 7.82** |
| Size x Growth | 0.08 | 0.04 | 0.03 |

**FIGURE 3:**
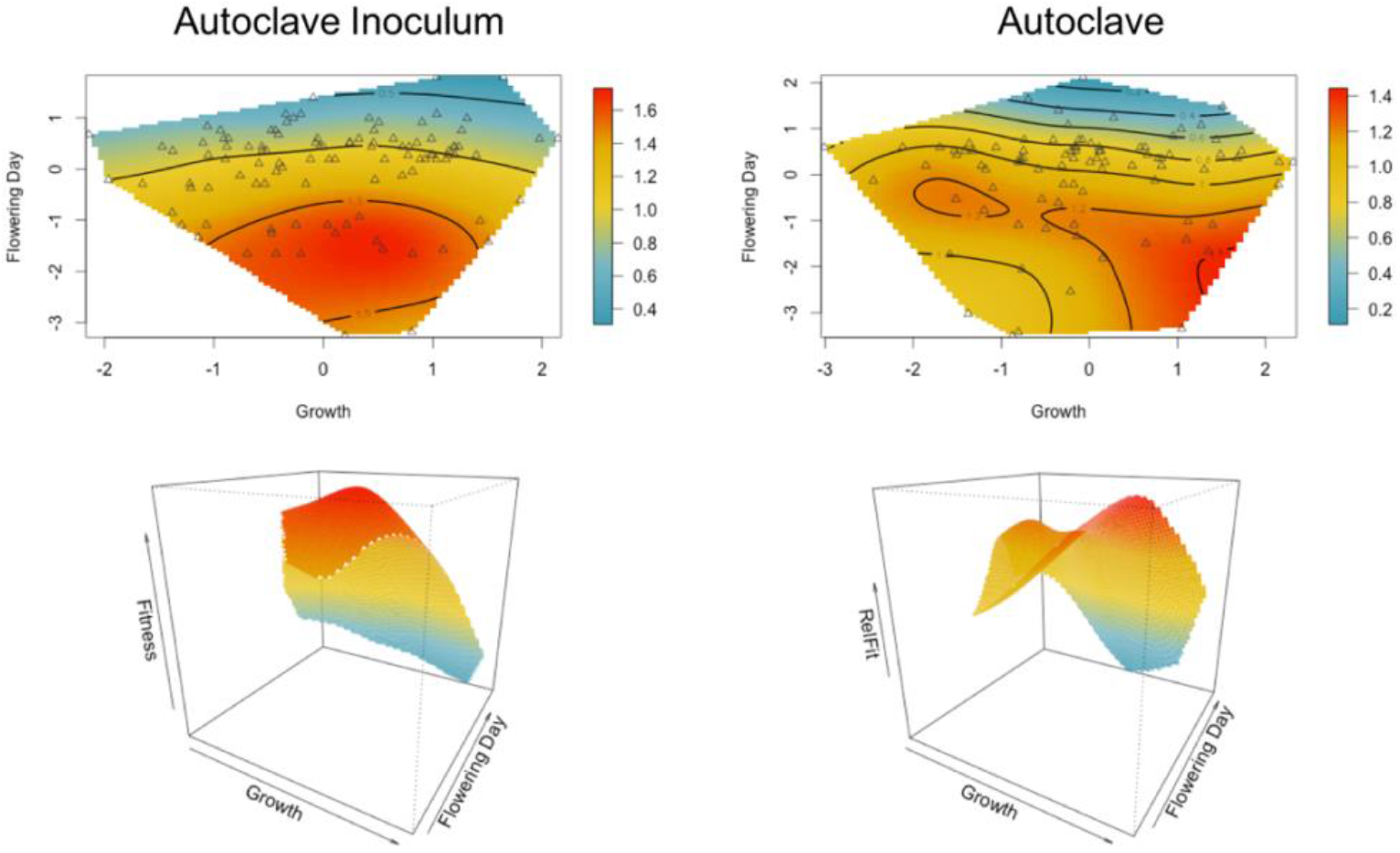
Correlational selection on *I. purpurea* flowering day and growth in a complex (autoclave + inoculum) and simple (autoclave) soil microbial community.

## DISCUSSION

The goal of the present work was to determine if the soil microbial community influences phenology and growth in the common morning glory, *I. purpurea*, and to determine if the soil community acts as an agent of selection on these important plant traits. We found evidence for both changes in plant phenotypes and altered patterns of selection as a result of soil microbial community manipulations. When grown in a complex soil microbial community, plants were larger and had more flowers than when exposed to a simple community. We also found that the soil microbial community influenced patterns of selection – while earlier flowering was favored in both the complex and simplified soil communities, selection for early flowering was stronger when plants were grown in the complex soil community. We likewise found evidence for changes in the pattern of correlational selection between the complex and simple soil microbial community environments. In the complex soil, there was evidence for one optimal growth/flowering time strategy whereas there were two peaks of high fitness in the simplified community – plants that grew fast and exhibited early flowering showed a fitness advantage as did slow growing plants that flowered later. This work adds to the growing literature investigating the role of the soil microbial community on plant phenology and other components of fitness. While on the whole we found broad similarities in plant responses between *I. purpurea* and the other (albeit few) species that have been investigated, we also uncovered important differences in the pattern of selection when soil microbial environments are altered. Below we discuss these similarities and differences in light of the accumulating evidence that the soil microbial community can influence plant evolutionary processes through its effect on plant phenology.

### Plant phenotypic changes as a result of microbial soil manipulation

We had *a priori* reasons to expect changes in flowering time in response to simplifying the soil microbial environment in *I. purpurea*. Previous research has shown that a simplified soil microbial community induces plant stress, which can lead to earlier flowering (Grime 1993). In support of this, Wagner et al. (2014) found that *B. stricta* flowered up to 2.8 days earlier in simplified compared to complex field soils. Further, increased growth rate and larger plant size has been associated with earlier flowering (Munguía-Rosas et al., 2011). If the complex soil microbial community in our experiment led to faster growth rates and larger plant size, individuals grown in the complex soil might flower earlier, leading to an observed delay of flowering in the simplified environment. Despite our broad expectations for a change in flowering time, however, we found no evidence that altering the soil microbial community by sterilizing the soil changed flowering time in the common morning glory, a result similar to that found in *B. rapa* (Lau and Lennon, 2011). Thus, our results do not allow us to draw general conclusions about how the microbial soil community may influence the timing of flowering, as it may be both species and experiment specific. Interestingly, Panke-Buisse et al. (2014) were able to alter flowering time of multiple plant species by first ‘priming’ the soil microbiome with later-flowering genotypes of *Arabidopsis*. The results of their work suggest that the absence of flowering time differences between environments in our experiment could simply be due to the absence of microbial species within the soil microbiome that exert an effect on flowering time.

We did find, however, that manipulation of the soil microbial community changed both plant size and the number of flowers produced in *I. purpurea*. Plants in the complex microbial soil community were larger and produced more flowers compared to *I. purpurea* grown in the simplified soil microbial community. These results are similar to that reported in *B. rapa* – plants grown in a complex environment were larger and produced more flowers than plants grown in a simplified microbe treatment (Lau and Lennon, 2011). While the presence of a diverse soil microbial community is known to positively impact plant productivity across a number of species (van der Heijden et al., 2008), how diverse soils may lead to increased plant biomass is not currently understood. Panke-Buisse et al. (2014) hypothesized that an increase in available nitrogen in the soil released by microorganisms could be responsible for increases in plant biomass. Another idea is that a greater diversity of interacting microbial species may be more effective at suppressing pathogenic microbes, such that more complex soils allow for greater and/or faster plant growth (Garbeva et al., 2004; Jousset et al., 2014; Hol et al., 2015).

### The soil microbial community as an agent of selection on phenology and growth

We found evidence for selection on flowering time in *I. purpurea* and that the soil microbial community acts as a selective agent on phenology in this species. In both soil environments we found flowering time to be under positive and stabilizing selection; plants that flowered earlier and/or at an intermediate time were at a fitness advantage regardless of soil microbial community environment. Strikingly, selection for early flowering was significantly stronger in the complex compared to the simplified soil environment – the change in soil environment altered the linear selection differential on flowering time by 55% between environments (Figure 2). This finding was opposite our expectations based on work in *B. rapa* where selection for earlier flowering was stronger when plants were grown in simplified rather than complex microbial communities (Lau and Lennon, 2011). Regardless, the change to the soil microbial community altered the strength of selection on flowering time, providing evidence that the soil microbial community can act as an agent of selection on flowering time in *I. purpurea*, as shown in *B. rapa* and *B. stricta* (Lau and Lennon, 2011; Wagner et al., 2014). Additionally, in our experiment, selection gradients and selection differentials were similar, indicating that the majority of selection on flowering time was direct and not indirect due to selection on correlated traits.

We likewise found evidence for positive linear selection on size in both soil environments, but no evidence that the strength or pattern of selection on plant size varied between simplified and complex soils. Thus, while larger plants were more fit in both soil types (as was found in *B. rapa*; Lau and Lennon, 2011), there was no indication that the soil microbial environment acted as an agent of selection on plant size in *I. purpurea*. Interestingly, although plant size was under selection, there was no indication that growth rate in *I. purpurea* in this experiment was under selection and correspondingly no evidence that selection on plant growth rate was altered by our microbial soil manipulations.

However, we found two different patterns of selection when we considered the potential for correlational selection between flowering time and growth rate. In the complex microbial soil environment, selection favored individuals that exhibited intermediate growth rates and an early- to mid-flowering time, whereas in the simplified soil community there was evidence for negative correlational selection between growth rate and flowering time. In this soil environment, there were two peaks of high fitness – one in which plants that grew fast and flowered early were at a fitness advantage, and another where slow growing and late-flowering plants exhibited a peak of high fitness. Thus, disruption to the soil community altered the dual combination of growth and flowering time phenotypes, meaning that the soil microbial community acts as an agent of selection on interactions between those traits. Although we do not know how the presence and abundance of microbial species may have changed over the course of this experiment, we suspect either the absence of particular microbial species or shifts in abundance of species over time led to differences in the pattern of selection on flowering time/growth combinations. It is likely that, for example, the peak of high fitness of fast growing, earlier flowering individuals was due to the stress of the simplified soil environment, whereas the peak of fitness corresponding to slow growing, later flowering individuals could have been due to later colonization of microbes from watering and other greenhouse conditions. This highlights an important caveat of the current study – we did not perform microbiome sequencing to identify the bacterial or fungal species present in the soil, and thus we cannot determine the underlying cause of the changed pattern of selection beyond that of broad manipulations of the soil microbiome. Regardless of the underlying microbial species (and changes to those species) that may be influencing our observed changes in the pattern of selection on *I. purpurea*, our soil preparation methods closely followed that of other studies that have previously established autoclaving as a means of simplifying the microbial community (Marschner and Rumberger, 2004; Berns et al., 2008; Lau and Lennon, 2011; Aschehoug et al., 2012; Panke-Buisse et al., 2014), and the observed phenotypic responses in this study (*i.e.*, lower plant biomass and fewer flowers produced in the simplified community) mirror that of this previous work.

Because we included a diverse set of genotypes in this study, we were able to examine the potential for genetic variation underlying phenology and plant size/growth traits in addition to potential selection on these traits. Similar to our previous work in *I. purpurea* (Chaney and Baucom, 2014), we identified maternal line and thus genetic variation in flowering time, indicating that flowering time can respond to selection imparted by changes to the soil community. We likewise identified genetic variation in plant size and total number of flowers produced across the twenty inbred accessions used, but not variation underlying growth rate. This suggests that although we find evidence of phenotypic correlational selection for two trait optima in the simplified soil community, we would expect the evolution of flowering time rather than evolution of two different life history strategies (*i.e.* either early flowering, fast growth, or late flowering, slow growth). Including maternal line variation in this study has thus allowed us to connect the ecological findings – *i.e.* the pattern of phenotypic selection – with expectations based on evolutionary principles. Also of note is the lack of genetic variation for flowering time and growth trait plasticity in our experiment, similar to Wagner et al. (2014), who found no evidence for adaptive plasticity in flowering time across different microbial treatments. Interestingly, their experiment examined the potential for adaptive plasticity in flowering time across different types of natural soils, whereas ours compared sterilized field soils to sterilized, re-inoculated field soils.

## CONCLUSION

Phenology plays an important role in plant evolutionary processes since the timing of flowering influences pollinator visitation and the potential for gene flow both within and among populations. The regulation of flowering time is primarily driven by abiotic factors, such as vernalization and photoperiod (Amasino, 2010), but stress, herbivory, genotype, and nutrient deficiencies can also influence phenology (Stanton et al., 2000; Franks et al., 2007; Jordan et al., 2015; Richardson et al., 2016). Here we add to the few available studies that consider the influence of the microbial community on plant flowering, and show that the soil microbial community acts as a selective agent on flowering phenology and combinations of phenology and growth in the common morning glory. The results of our work suggest that flowering time evolution should respond more rapidly in environments with complex soil communities compared to simplified soil communities, a finding that if common and consistent in natural settings would suggest that flowering time evolution may be slowed in microbe-poor soils. However, more studies of a diverse range of taxa should be undertaken before the conclusion is drawn, especially given that the opposite pattern was previously shown in *B. rapa*, where stronger linear selection on flowering time was identified in the simple rather than the complex microbial environment (Lau and Lennon, 2011). Examining changes to plant flowering time in light of environmental influences – including that of the soil microbiota – can provide useful tools for understanding how plants may alter their life cycles to cope with environmental heterogeneity and episodes of plant stress. To date, very few studies have considered the interaction of microbial community changes and components of the abiotic environment, and thus more work remains in dissecting the influence of complex and interacting forces on flowering time evolution across diverse plant species.

## Acknowledgments

The authors would like to thank Laurie Elliott Nichols and Deborah LeGendgre for their work on this experiment.

## Data Accessibility Statement

Data for this study will be made available via Dryad post-acceptance.

## SUPPORTING INFORMATION

Additional Supporting Information may be found online in the Supporting Information section at the end of the article.

## APPENDIX S1

Selection differentials (univariate selection analysis; *S*) for *Ipomoea purpurea* plant traits. Shown are selection values and model *F*-values. Significant effects are indicated with bold font and asterisks: \*\*\**P* < 0.001.

## APPENDIX S2

Selection gradients (multivariate selection analysis) for *Ipomoea purpurea* plant traits. Shown are linear (beta; β) and quadratic (gamma; γ) values. Linear coefficients were determined in each treatment from the first-order model only, whereas the second-order coefficients were determined from the full model with the linear, squared and cross-product terms. Quadratic regression coefficients were converted to selection gradients by doubling them. Significant effects are indicated with asterisks: \*\*\**P* < 0.001.

