## Appendix for "The soil microbial community alters patterns of selection on flowering time and fitness related traits in *Ipomoea purpurea*"

**APPENDIX S1:** Selection differentials (univariate selection analysis; *S*) for *Ipomoea purpurea* plant traits. Shown are selection values and model *F*-values. Significant effects are indicated with bold font and asterisks: ***$P < 0.001$.

| | Selection (*S*) | ***F*-value** |
|---|---|---|
| Flowering day | **-0.22** | 36.78*** |
| Size | **0.14** | 14.96*** |
| Growth | -0.03 | 0.45 |

**APPENDIX S2:** Selection gradients (multivariate selection analysis) for *Ipomoea purpurea* plant traits. Shown are linear (beta; β) and quadratic (gamma; γ) values. Linear coefficients were determined in each treatment from the first-order model only, whereas the second-order coefficients were determined from the full model with the linear, squared and cross-product terms. Quadratic regression coefficients were converted to selection gradients by doubling them. Significant effects are indicated with asterisks: ***$P < 0.001$.

| | **linear (β)** | **quadratic (γ)** |
|---|---|---|
| Flowering day | -0.20*** | -0.26*** |
| Size | 0.18*** | -0.01 |
| Growth | -0.10*** | -0.02 |
| Flowering day x size | | -0.08 |
| Flowering day x growth | | -0.04 |
| Size x Growth | | 0.03 |
